# Multimodal single-cell chromatin analysis with Signac

**DOI:** 10.1101/2020.11.09.373613

**Authors:** Tim Stuart, Avi Srivastava, Caleb Lareau, Rahul Satija

## Abstract

The recent development of experimental methods for measuring chromatin state at single-cell resolution has created a need for computational tools capable of analyzing these datasets. Here we developed Signac, a framework for the analysis of single-cell chromatin data, as an extension of the Seurat R toolkit for single-cell multimodal analysis. Signac enables an end-to-end analysis of single-cell chromatin data, including peak calling, quantification, quality control, dimension reduction, clustering, integration with single-cell gene expression datasets, DNA motif analysis, and interactive visualization. Furthermore, Signac facilitates the analysis of multimodal single-cell chromatin data, including datasets that co-assay DNA accessibility with gene expression, protein abundance, and mitochondrial genotype. We demonstrate scaling of the Signac framework to datasets containing over 700,000 cells.

**Availability:** Installation instructions, documentation, and tutorials are available at: https://satijalab.org/signac/

## Introduction

Several technologies are now available for measuring aspects of chromatin state at single-cell resolution, particularly single-cell (sc) ATAC-seq and scCUT&Tag [Ai et al., 2019, Buenrostro et al., 2015, Carter et al., 2019, Cusanovich et al., 2015, Kaya-Okur et al., 2019, Lareau et al., 2019, Luo et al., 2018, Satpathy et al., 2019, Smallwood et al., 2014]. The development of these new technologies has created a need for computational tools to analyze single-cell chromatin data. While the analysis of these datasets presents some unique challenges in comparison to more established single-cell methods like scRNA-seq, many analysis steps are shared. These include nonlinear dimension reduction, cell clustering, identifying differentially active features between groups of cells, and visualizing cells in reduced dimension space. Alongside these common tasks, the analysis of single-cell chromatin data present opportunities for several more specialized analysis tasks. These include identifying DNA sequence features (motifs or variants) that are enriched in different sets of cells, specialized feature weighting and linear dimension reduction methods, and genome browser-style data visualization.

Furthermore, new technologies now enable the co-assay of multiple cellular modalities in single cells, including DNA accessibility alongside mRNA abundance [Cao et al., 2018, Chen et al., 2019, Clark et al., 2018, Ludwig et al., 2019, Lareau et al., 2020, Zhu et al., 2019, Xing et al., 2020, Liu et al., 2019, Ma et al., 2020], protein abundance [Mimitou et al., 2020, Fiskin et al., 2020, Swanson et al., 2020], CRISPR guide RNAs [Rubin et al., 2019, Pierce et al., 2020], or spatial position [Thornton et al., 2019]. These datasets present unique opportunities to learn the relationships between cellular modalities [Stuart and Satija, 2019], and will be especially powerful in deciphering the regulatory roles of noncoding DNA sequences. The analysis of these datasets is challenging without software designed to facilitate a multimodal analysis, and an ideal computational solution would facilitate an integrative analysis of multimodal single-cell data encompassing gene expression, chromatin state, and other modalities including cell lineage, protein expression, or spatial position in a single framework. However, existing tools for the analysis of single-cell chromatin data were designed for the analysis of unimodal single-cell datasets [Danese et al., 2019, Fang et al., 2019, Granja et al., 2020].

Here we developed Signac, a framework for the analysis of single-cell chromatin data, as an extension of the Seurat toolkit [Satija et al., 2015, Butler et al., 2018, Stuart et al., 2019]. By building on the existing Seurat package, Signac allows for the analysis of multimodal single-cell data by accessing the extensive existing computational methods available in the Seurat package and in other packages that interface with the Seurat object. Signac enables the end-to-end analysis of chromatin data and includes functionality for diverse analysis tasks, including: identifying cells from background non-cell-containing barcodes, calling peaks, quantifying counts in genomic regions, quality control filtering of cells, dimension reduction, clustering, integration with single-cell gene expression data, interactive genome browser-style data visualization, finding differentially accessible peaks, finding enriched DNA sequence motifs, transcription factor footprinting, and linking peaks to potential regulatory target genes (Figure 1A). Furthermore, Signac provides a framework for the identification of mitochondrial genome variants from single-cell DNA accessibility experiments, enabling a joint analysis of clonal relationships and DNA accessibility in single cells [Lareau et al., 2020, Ludwig et al., 2019, Xu et al., 2019].

**Figure 1.**
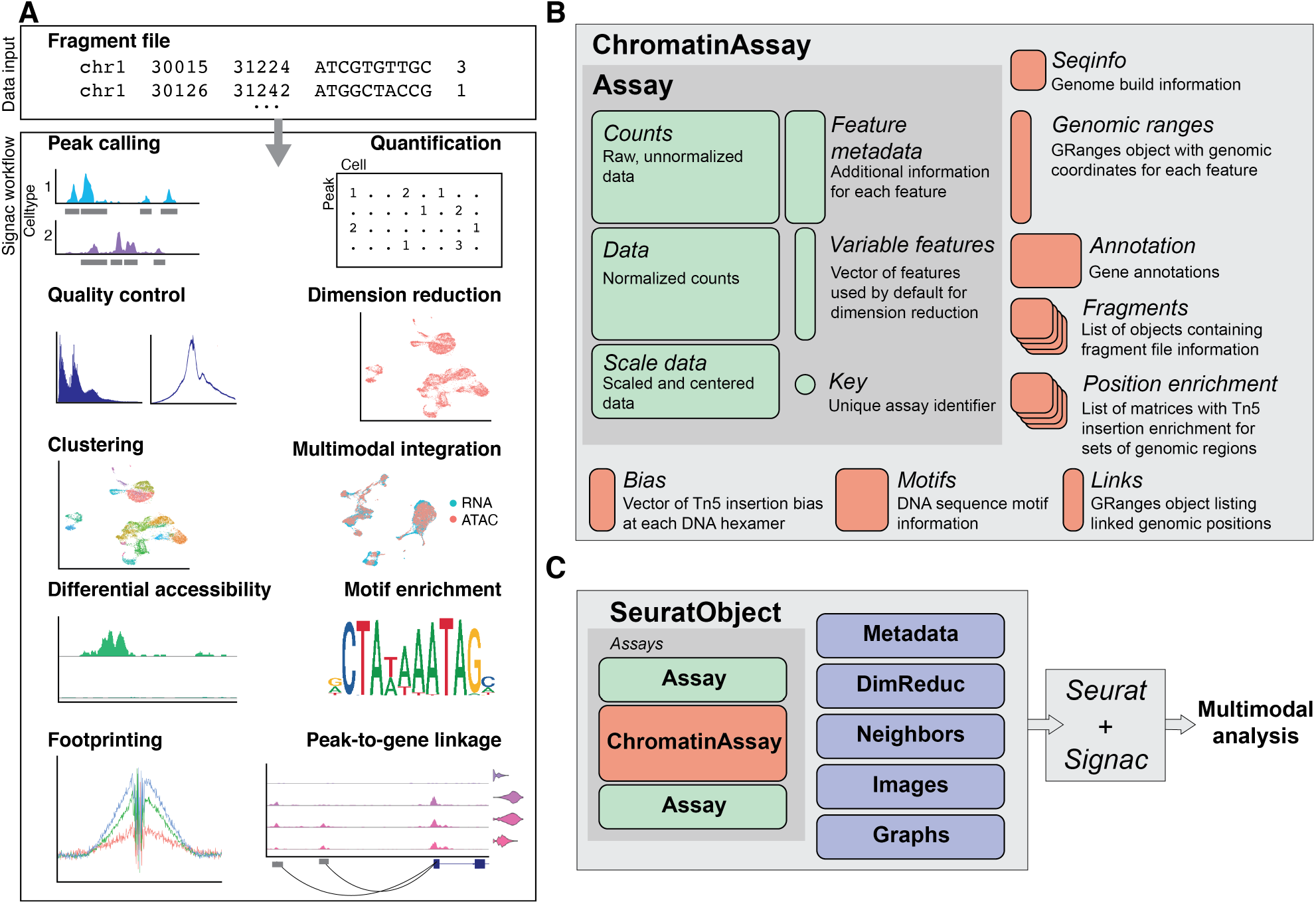
Single-cell chromatin analysis workflow with Signac. (A) Overview of the key steps comprising the analysis of a single-cell chromatin data with Signac. All analysis tasks can be completed with one or multiple fragment files as input. (B) Design of a custom *Assay* for single-cell chromatin data. We extended the existing Seurat *Assay* to add the ability to store data required for the analysis of single-cell chromatin datasets. (C) Extended *ChromatinAssay* objects can be stored side-by-side with standard *Assay* objects in Seurat to enable the analysis of multimodal single-cell data.

## Results

### Package design

We aimed to create an extensible framework for single-cell chromatin data analysis that builds on existing tools used in the single-cell, genomics, and R-language communities. We developed an R toolkit for the analysis and visualization of single-cell chromatin data as an extension of our existing Seurat package, designed for the analysis of multimodal single-cell data [Butler et al., 2018, Stuart et al., 2019, Hao et al., 2020]. The Seurat package uses the *Seurat* object as its central data structure. The *Seurat* object is composed of any number of *Assay* objects containing data for single cells. The *Assay* object was originally designed for the analysis of single-cell gene expression data, and allows for storage and retrieval of raw and processed single-cell measurements and metadata associated with each measured feature. To facilitate the analysis of single-cell chromatin data within the Seurat framework, we developed an extended *ChromatinAssay* object class (Figure 1B). The *ChromatinAssay* extends the Seurat *Assay* to allow for the storage and retrieval of information needed for the analysis of single-cell chromatin data, including genomic ranges associated with each feature in the experiment, gene annotations, genome build information, DNA motif information, and on-disk storage of single-cell data as tabix-indexed fragment files [Li, 2011]. We leveraged existing data structures used in the Bioconductor community [Gentleman et al., 2004, Huber et al., 2015] for interacting with genomic ranges and position-indexed genomic data files [Arora et al., 2020, Lawrence et al., 2013, Li, 2011, Morgan et al., 2020]. Furthermore, we provide flexible parallelization strategies through the future R package [Bengtsson, 2020], and extensible plotting with the ggplot2 package [Wickham, 2016]. Crucially, the extended *ChromatinAssay* can be stored in a Seurat object side-by-side with standard Seurat *Assay*-class objects to facilitate the analysis of multimodal single-cell data (Figure 1C).

### Analysis of multimodal human PBMC data

To demonstrate the core functionality of the Signac package we analysed a publicly available dataset that jointly profiled mRNA abundance and DNA accessibility in single human pe-ripheral blood mononuclear cells (PBMCs), generated by 10x Genomics. We computed per-cell quality control (QC) metrics using the DNA accessibility assay, including the strength of the nucleosome banding pattern and transcriptional start site (TSS) enrichment score (Figure 2A; see Methods) and removed cells that were outliers for these QC metrics. Next, we processed the gene expression assay by normalizing the RNA counts with SCTransform and Seurat [Hafemeister and Satija, 2019, Stuart et al., 2019]. Patterns of DNA accessibility can be difficult to interpret alone, as much less is currently known about the patterns of cell-type-specific DNA accessibility and the function of noncoding DNA elements than is known about the function and cell-type specificity of protein-coding gene expression. We therefore chose to annotate cell types by mapping the cells to an annotated multimodal PBMC reference dataset, using the gene expression assay [Hao et al., 2020]. This revealed 23 different cell types present in the dataset, including rare populations such as gamma delta T cells.

**Figure 2.**
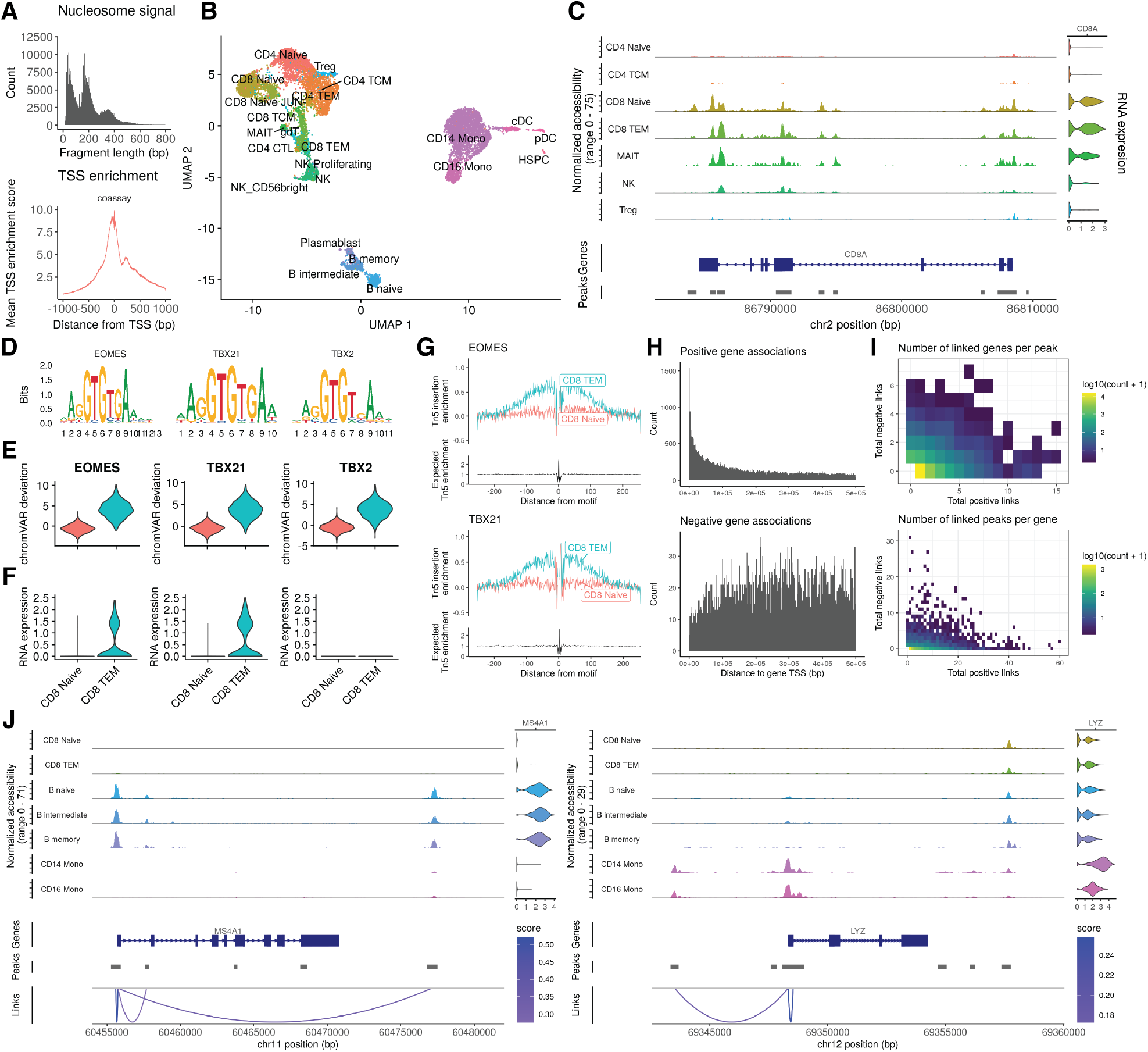
Integrative single-cell analysis of gene expression and DNA accessibility in human PBMCs. (A) Quality control metrics for single-cell chromatin data. Nucleosome signal and transcriptional start site (TSS) enrichment score metrics can be used to identify low-quality cells for removal prior to downstream analysis. (B) Joint UMAP representation of the multimodal human PBMC dataset, with cells annotated by predicted cell type. (C) Genome browser visualization of combined DNA accessibility and gene expression data at the *CD8A* locus, displaying differential DNA accessibility among naive and effector CD8+ T cells. (D) DNA sequence motifs for top overrepresented TF motifs between CD8+ effector and naive T cells. (E) chromVAR [Schep et al., 2017] deviations for top enriched DNA sequence motifs (*EOMES, TBX21, TBX2*) for CD8+ effector (CD8 TEM) and naive CD8+ (CD8 Naive) T cells. (F) RNA expression for *EOMES, TBX21*, and *TBX2* genes in CD8+ effector and naive T cells. (G) TF footprinting analysis for *EOMES* and *TBX21* motifs sites. (H) Distances from peak to linked gene TSS, for positive and negative coefficient peak-gene links. (I) Total number of positive-coefficient and negative-cofficent peak-gene links for each linked gene (top) and peak (bottom). (J) Representative example peak-gene links for key immune genes.

The analysis of chromatin datasets can be highly dependent on accurate peak calling, and this challenge is compounded in single-cell assays where peaks specific to rare populations are sometimes missed when calling peaks on the whole cell population. To address this problem, we identified peaks using MACS2 [Zhang et al., 2008] for each annotated cell type separately and combined the individual peak calls into a unified peak set using Signac. Indeed, peaks specific to rare cell populations were often missed when calling peaks on the whole dataset (Figure S1 A, B). We further compared the MACS2 cell-type-specific peak calls with the peak calls produced by 10x Cellranger, commonly used for the analysis of scATAC-seq data, and found 15,777 cases where a Cellranger peak merged distinct MACS2 peaks into a single region, whereas there were no cases where a MACS2 peak overlapped multiple Cellranger peaks (Figure S1C). This revealed a bias in Cellranger for aberrant merging of multiple distinct peaks into a single region, and highlights the importance of accurate cell-type-specific peak calling methods in the analysis of single-cell chromatin datasets.

We next reduced the dimensionality of the DNA accessibility assay by latent semantic indexing (LSI) [Cusanovich et al., 2015, Deerwester et al., 1990], and reduced the dimensionality of the gene expression assay by principal component analysis (PCA). To construct a low-dimensional representation of the cells that reflected both data modalities we applied the recently developed weighted nearest neighbor (WNN) methods to construct a joint neighbor graph representing both data modalities [Hao et al., 2020], and used this graph to construct a joint UMAP visualization [McInnes and Healy, 2018] (Figure 2B). This revealed the diversity of cell states present in the dataset, and highlighted the differing power of the two modalities to separate the different cell states present in the dataset. For example, regulatory T cells (Treg) were able to be better separated using the gene expression data, while CD4+ naive and CD8+ naive T cells were better separated using the DNA accessibility data (Figure S2).

To explore the differences in chromatin landscapes between cell types in the PBMC dataset, we identified ATAC-seq peaks open in CD8+ effector T cells relative to CD8+ naive T cells, revealing many regions of open chromatin that were specific to the CD8+ effector cells (Figure 2C). To identify transcription factors (TFs) that may be implicated in regulating these cells, we searched for overrepresented DNA sequence motifs in the set of CD8+ effector cell-specific peaks (see Methods). This revealed a strong overrepresentation of *EOMES, TBX21*, and *TBX2* TF binding motifs. However, the motifs for each of these TFs are nearly identical (Figure 2D) and displayed the same patterns of accessibility among the cells (Figure 2E), making it difficult to correctly identify the TF involved in binding these motifs in effector T cells. To identify the putative regulatory TFs, we examined the gene expression data in these cells. While *EOMES* and *TBX21* were both expressed in T cells, *TBX2* was not detected (Figure 2F). This indicated that *EOMES* and *TBX21* likely regulate these sites [Pearce et al., 2003], rather than *TBX2*, and highlights the ability of combined gene expression and DNA accessibility data to improve the identification of TFs involved in regulating different cell states. We further examined the enrichment of Tn5 integration events surrounding *EOMES* and *TBX21* motifs sites by performing TF footprinting [Corces et al., 2018], revealing a strong enrichment of integration events flanking the TF motif in CD8+ effector cells compared to CD8+ naive cells (Figure 2G).

The measurement of both gene expression and DNA accessibility in the same cell creates an opportunity to link noncoding DNA elements to their potential regulatory targets through the correlation between DNA accessibility and the expression of a nearby gene. Indeed, many past studies that measured both DNA accessibility and gene expression in the same cell have performed a peak-to-gene linkage analysis using regression models [Cao et al., 2018, Zhu et al., 2019, Ma et al., 2020]. We implemented a peak-to-gene linkage method in Signac based on recently described methods [Ma et al., 2020]. Briefly, we computed the Pearson correlation between the expression of a gene and the accessibility of each peak within 500 kb of the gene transcriptional start site (TSS), and compared this value with the expected value given the GC content, overall accessibility, and length of the peak (see Methods). Applying this linkage method to all expressed genes in the PBMC dataset revealed a set of 37,424 peak-gene links across the genome. The vast majority of these links displayed a positive relationship between the accessibility of the peak and expression of the linked gene, with 89% of links having a positive correlation coefficient. Although links were enriched in close proximity to the gene TSS, we also observed a substantial number of long-range putative regulatory relationships, with 58% of links spanning a distance of >100 kb from the gene TSS (Figure 2H). Linked genes were on average linked to ∼6 peaks (mean = 6.37, standard deviation = 7.09), while linked peaks were linked to ∼1 gene on average (mean = 1.57, standard deviation = 1.26) (Figure 2I).

Cell-type-specific immune genes appeared to form accurate links with nearby peaks accessible in the same cell types expressing the genes (Figure 2J). We sought to systematically assess the accuracy of the peak-gene links identified by examining a set of expression quantitative trait loci (eQTL) fine-mapped variants for whole blood, produced by the GTEx consortium [GTEx Consortium, 2020]. For linked peaks that overlapped a fine-mapped eQTL variant, the eQTL variant was associated with the same gene as the peak in 52.6% of cases, while 13.4% was expected by random chance. The systematic identification of putative regulatory targets for any open chromatin region in the genome using multimodal single-cell datasets has the potential to enable a more accurate assignment of trait- or disease-associated noncoding variants to a gene likely to be impacted by the variant. Furthermore, the identification of distal regulatory elements for each gene in the genome will enable these sequences to be included in predictive models of gene expression, enabling the accuracy of these models to be improved [Agarwal and Shendure, 2020, Kelley et al., 2018].

### Comparison of LSI dimension reduction methods

LSI was originally developed for natural language processing [Deerwester et al., 1990], and uses a term frequency-inverse document frequency (TF-IDF) weighting scheme to weight features according to their frequency in a document and their frequency across all documents in a text corpus. LSI has since been applied for the analysis of single-cell chromatin data, where a cell is analogous to a document and a term analogous to a peak or genomic region [Cusanovich et al., 2015]. The most popular TF-IDF method applied to single-cell chromatin data computes the term frequency as *TF* = *C*_*i j*_*/F*_*j*_ where *C*_*i j*_ is the total number of counts for peak *i* in cell *j* and *F*_*j*_ is the total number of counts for cell *j*. The inverse document frequency is typically computed as *IDF* = *log*(1 + *N/n*_*i*_) where *N* is the total number of cells in the dataset and *n*_*i*_ is the total number counts for peak *i* across all cells. The TF-IDF matrix is then computed as *TF*× *IDF*. We found that, when applied to scATAC-seq data, this implementation often results in nonzero values in the TF-IDF matrix having low variance and a mean very close to zero, and a poor ability to discriminate between cell types. We developed a simple modification to the TF-IDF weighting scheme that improves the results of LSI when applied to single-cell chromatin data (Figure 3A). In our modified method we compute the inverse document frequency as *IDF* = *N/n*_*i*_, and TF-IDF as *log*(1 + (*TF* × *IDF*) × 10^4^).

**Figure 3.**
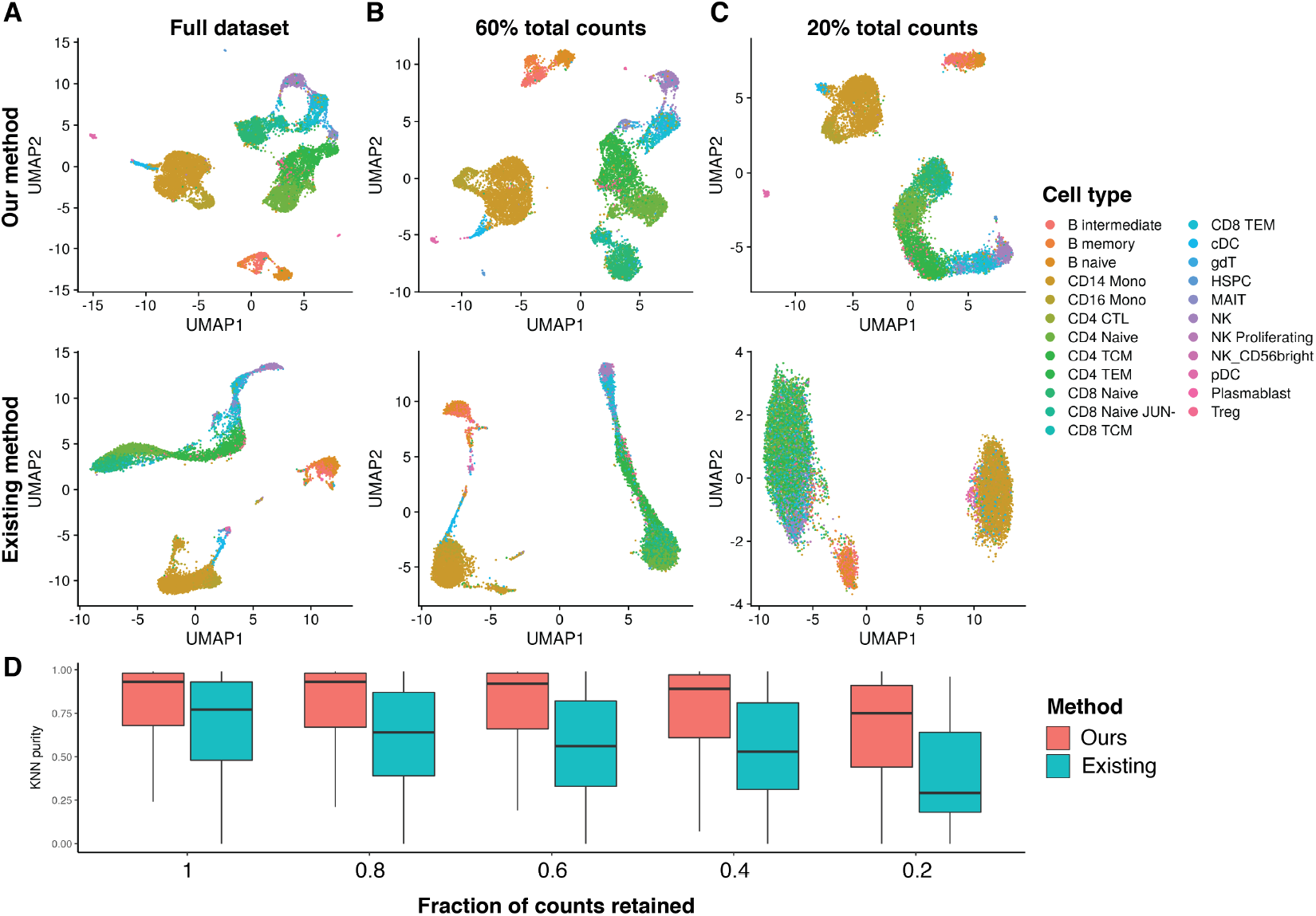
Impact of different TF-IDF methods on LSI performance. (A) LSI and UMAP performed on the PBMC dataset at full depth, and downsampled to (B) 60% of the total counts, or (C) 20% total counts. (D) K-nearest neighbor purity metric for each TF-IDF method and for each downsampled dataset.

To test the performance of our modified TF-IDF method, we downsampled the total counts for the multimodal PBMC dataset and performed LSI and UMAP using the original TF-IDF method [Cusanovich et al., 2015] and our modified method. When using our modified TF-IDF method, cell types were able to be separated even when using 20% of the counts of the full dataset, whereas this was not possible when using the original TF-IDF method (Figure 3B, C). We further assessed the preservation of local cell neighborhoods in each downsampled dataset by computing the fraction of k-nearest neighbors (k=100) for each cell belonging to the same cell type as the query cell (KNN purity), with cell types annotated using the independent gene expression assay. This revealed a gradual decline in local structure preservation as fewer counts were retained from the original dataset, with a greater decline seen when using the original TF-IDF method (Figure 3D). These results indicate that LSI, when applied with the right TF-IDF method, can be a powerful dimension reduction technique for single-cell DNA accessibility data. Furthermore, LSI is scalable to large numbers of cells as it retains the data sparsity (zero counts remain as zero after applying TF-IDF). This is not the case for other methods such as the Jaccard similarity [Fang et al., 2019]. LSI also uses the singular value decomposition (SVD), for which there are highly optimized, fast algorithms that are able to run on sparse matrices [Baglama et al., 2019, Baglama and Reichel, 2005].

### Joint analysis of DNA accessibility and mitochondrial genotype

New technologies capable of measuring chromatin state alongside other data modalities at single-cell resolution are now being rapidly developed. These include the development of assays that measure DNA accessibility data alongside mitochondrial genome sequence [Lareau et al., 2020, Ludwig et al., 2019, Xu et al., 2019]. As the mitochondrial genome mutates at a much higher rate than the nuclear genome, and mitochondrial mutations are inherited over cell divisions, measuring mitochondrial genome sequences in single cells can be informative in reconstructing clonal cell relationships [Ludwig et al., 2019, Xu et al., 2019, Lareau et al., 2020]. These experiments therefore provide an opportunity to study DNA accessibility differences between or within clonal groups of cells. To facilitate a joint analysis of these datasets, we developed computational methods to enable the identification of informative mitochondrial variants, the calculation of mitochondrial variant allele frequencies, and clonal cell clustering within the Signac framework.

We analyzed a recently published single-cell DNA accessibility and mitochondrial genome sequence coassay dataset from a patient with a colorectal cancer (CRC) tumor [Lareau et al., 2020]. We first performed QC, dimension reduction, and clustering on the DNA accessibility assay, and annotated the major cell types present in the dataset based on the DNA accessibility at key marker genes (Figure 4A). This revealed five major clusters present in the dataset encompassing tumor-derived epithelial cells, basophils, myeloid cells, and T cells, as previously reported [Lareau et al., 2020]. To identify the clonal relationships between cells in the CRC dataset, we identified highly variable mitochondrial genome positions among the cells by computing the variance-mean ratio and the Pearson correlation between strand coverage (Figure 4B). Visualization of per-cell allele frequencies (fraction heteroplasmy) for these variants in the two-dimensional UMAP space computed using the DNA accessibility assay revealed the variant 16147C>T present at nearly 100% frequency in the tumor-derived epithelial cells, while other variants were shared across different immune cell types (Figure 4C). We further identified cell clones by clustering the allele frequency data, revealing 10 distinct clones (Figure 4D). Clones 1, 2, and 4 were highly specific to the epithelial cells, whereas other clones were dispersed more evenly across the different immune cell types, indicating that those immune cells likely originated from a common hematopoietic progenitor. We further identified differential DNA accessibility peaks between the three different epithelial cell clones, highlighting the ability of the additional clonotype data to aid in identifying additional sources of chromatin state heterogeneity within a cell type (Figure 4E).

**Figure 4.**
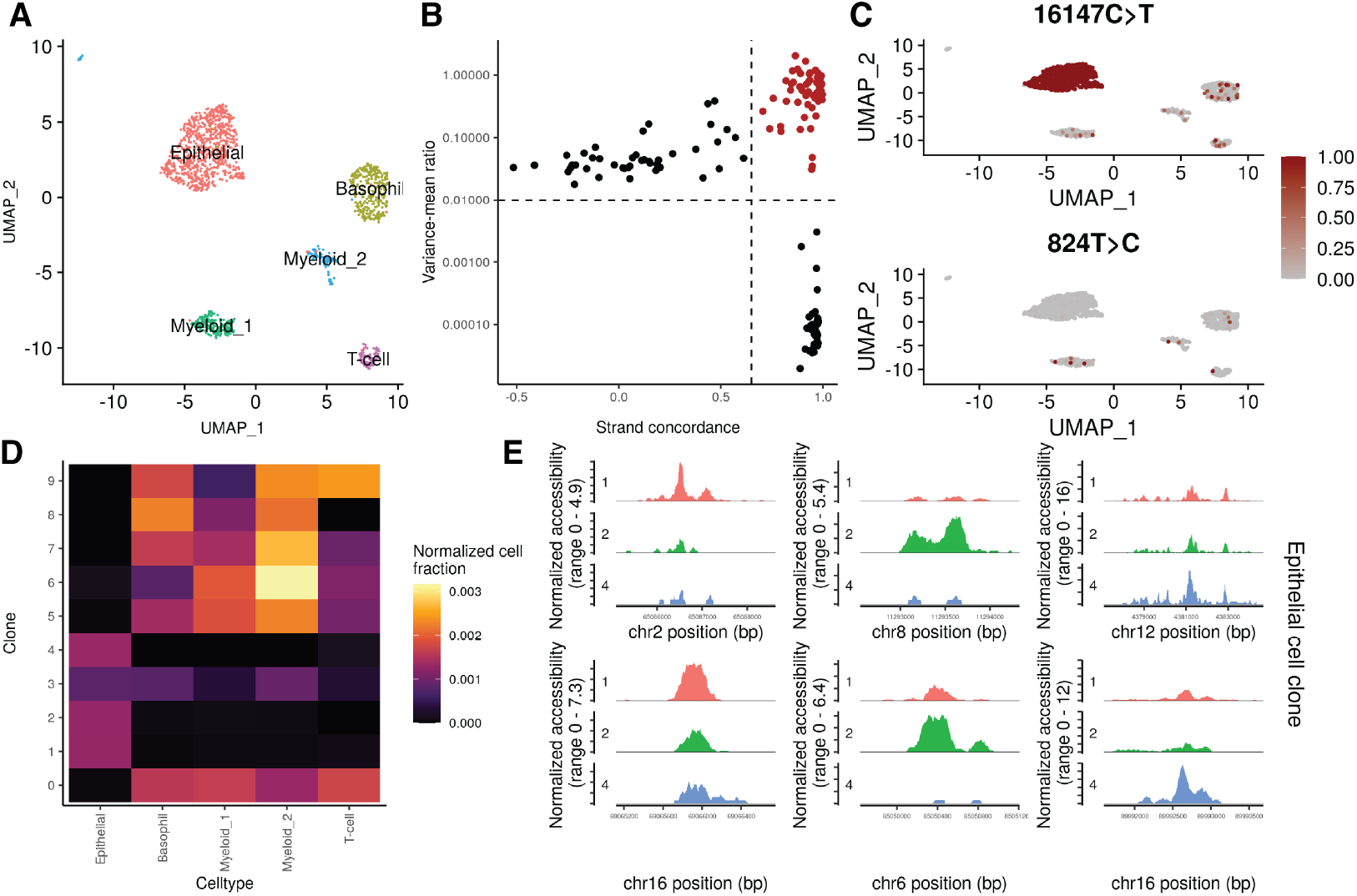
Joint analysis of mitochondrial genotypes and DNA accessibility in single cells. (A) UMAP plot for cells from a colorectal cancer patient tumor profiled by scATAC-seq, with the major cell types annotated. (B) Variance-mean ratio versus strand concordance (Pearson correlation between strand coverage) for mitochondrial genome variants. High-confidence, highly variable mitochondrial genome sites are shown in red. (C) Per-cell allele frequencies (fraction heteroplasmy) for two representative mitochondrial genome variants used to identify cell clones. (D) Fraction of cells belonging to each clone that were assigned to each cell type, normalized for the total number of cells beloning to each cell type. (E) Differentially accessible regions of the nuclear genome between epithelial cell clones.

### Scalable analysis of single-cell chromatin data

Methods are now available that enable the generation of very large scATAC-seq datasets [Lareau et al., 2019]. This presents opportunities to deeply characterize the chromatin state of tissues at single-cell resolution, but also raises the need for computational tools that similarly scale to large cell numbers. To demonstrate the scalability of Signac to datasets of this size, we re-analyzed a recently published scATAC-seq dataset from several regions of the adult mouse brain generated by the Brain Initiative Cell Census Network (BICCN) [Li et al., 2020]. After removing low-quality cells defined by low nucleosome signal and TSS enrichment QC metrics, this dataset contained 733,526 cells from 45 different brain regions.

We processed the entire dataset using a similar analysis workflow to that shown above for PBMCs, including the quantification of fragment counts per cell for each peak, computing per-cell QC metrics, and dimension reduction (Figure 5A). This revealed the diversity of cell types present in the mouse brain, as shown previously [Li et al., 2020]. To explore how the runtime and memory usage of different analysis steps scaled with different cell numbers, we downsampled the total number of cells in the BICCN dataset from 733,526 (the full dataset) down to 50,000 cells and re-ran the quantification (FeatureMatrix function), quality control (NucleosomeSignal and TSSEnrichment functions), and dimension reduction (RunTFIDF and RunSVD functions) steps (Figure 5B-F). For the FeatureMatrix step, which can be run in parallel, we also tested with 1, 2, 4 or 8 cores (Figure 5B). These results revealed a general trend of approximately linear increases in runtime and memory requirements for increasing dataset sizes, and provides a valuable benchmark resource for those planning experiments and estimating the time and resources required to analyze single-cell chromatin datasets with the Signac package.

**Figure 5.**
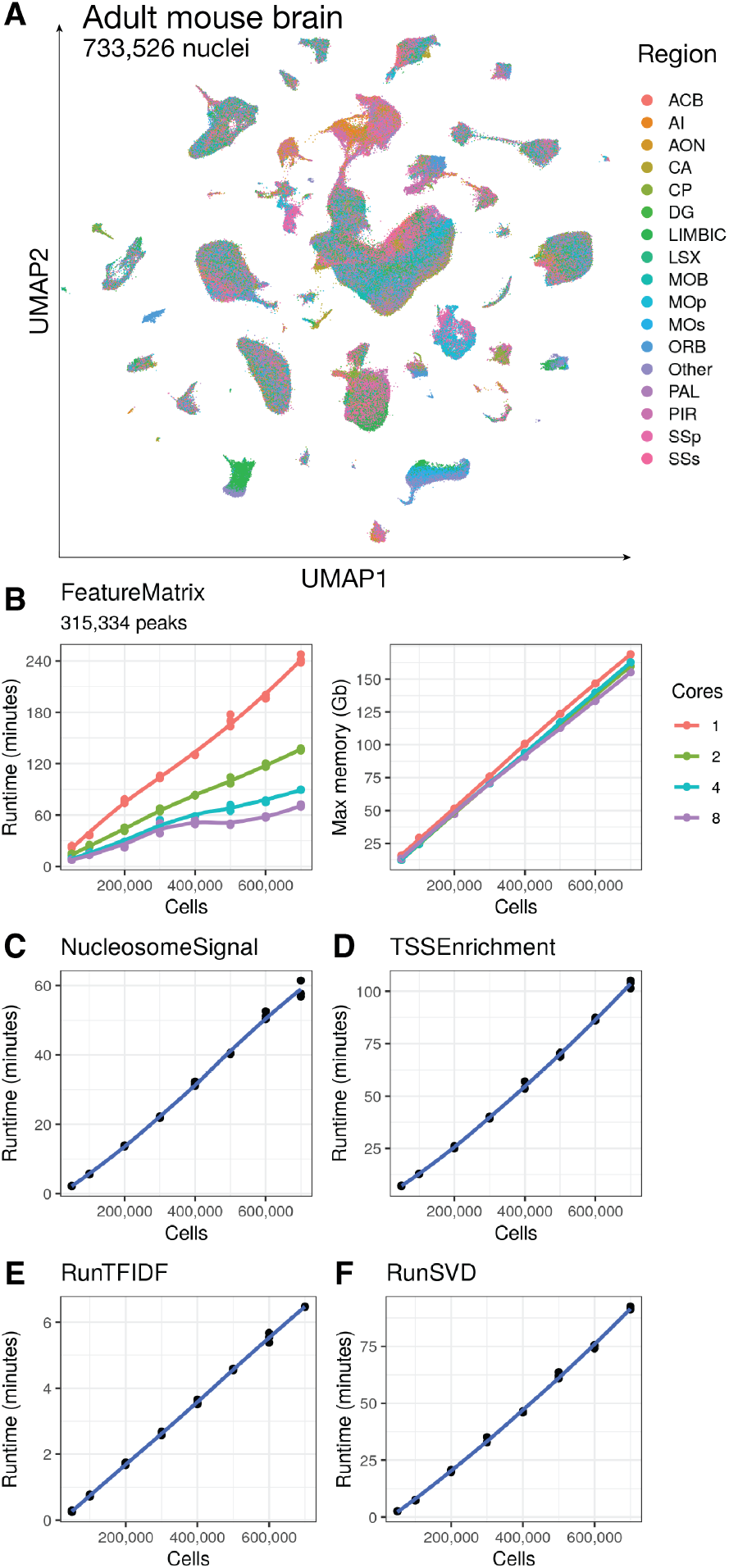
Scalable analysis of scATAC-seq data. (A) UMAP projection of the full BICCN mouse brain scATAC-seq dataset, with cells colored by the region of the mouse brain that they were sampled from. Runtime and peak memory usage for running (B) FeatureMatrix, (C) NucleosomeSignal, (D) TSSEnrichment, (E) RunTFIDF, and (F) RunSVD for varying numbers of cells.

## Discussion

As experimental methods for measuring aspects of chromatin state at single-cell resolution continue to be developed and improved, the parallel development of computational tools designed to analyse these datasets becomes increasingly important. Here, we developed Signac for the analysis of single-cell chromatin data, building on our existing Seurat toolkit [Butler et al., 2018, Stuart et al., 2019, Hao et al., 2020], and demonstrated running key analysis steps using Signac for the analysis of both unimodal and multimodal single-cell chromatin datasets. These analysis steps can be scaled to datasets containing >700,000 cells, and the scalability of these methods will become particularly important as large-scale cell atlas projects are completed [Li et al., 2020]. We further developed a simple modification to the popular LSI dimension reduction method that improved the performance of LSI when applied to single-cell chromatin data, particularly for datasets with low sensitivity. Furthermore, Signac enables running other tools developed by the community for the analysis of single-cell chromatin data, including chromVAR for estimating DNA motif variability between cells [Schep et al., 2017], Monocle for building pseudotime trajectories [Cao et al., 2019], Cicero for finding co-accessible networks of peaks [Pliner et al., 2018], and Harmony for performing dataset integration [Korsunsky et al., 2019]. As additional experimental methods for measuring multiple aspects of cell state are developed, a major challenge is to analyze these diverse datasets together in a consistent framework to learn how different modalities influence one another. The Seurat framework, via the extensible *Assay* class, is an appealing solution for the analysis of multimodal single-cell data, and we envision future computation methods will further build on the Seurat and Signac frameworks to jointly analyze multimodal single-cell datasets. A major challenge currently facing biology is understanding how the genome encodes the organism [Brenner, 2010]. Developing a deep understanding of how genes are regulated by noncoding DNA elements would greatly improve our ability to predict the effect of mutations, and to predict the target genes for trait-associated non-coding loci. A joint analysis multimodal single-cell chromatin and gene expression data hold great promise in furthering these goals, and the analytical framework presented here will be a valuable component in deciphering these gene regulatory relationships.

## Code and data availability

Signac is available on CRAN (https://cloud.r-project.org/package=Signac) and on GitHub (https://github.com/timoast/signac), with documentation and tutorials available at https://satijalab.org/signac/. All code used in this paper is available on GitHub: https://github.com/timoast/signac-paper. All data used in the paper is publicly available. The PBMC multiomic dataset is available from 10x Genomics: https://support.10xgenomics.com/single-cell-multiome-atac-gex/datasets/1.0.0/pbmc_granulocyte_sorted_10k. Data from the Brain Initiative Cell Census Network is available from the Neuroscience Multi-omic Archive (NeMO): https://nemoarchive.org/. Data for the colorectal cancer patient sample is available on NCBI GEO (GSE148509) and Zenodo (https://zenodo.org/record/3977808).

## Methods

Signac 1.1.0 was used for all analyses and is available on CRAN (https://cloud.r-project.org/package=Signac) and GitHub (https://github.com/timoast/signac/). R version 4.0.3 was used for all analyses, with standard BLAS and LAPACK libraries linked, running on Ubuntu 18.04.4 LTS with Intel Xeon W-2135 CPUs at 3.70GHz.

## Data structures

We extended the Seurat *Assay* class via the R-language class inheritance framework to create the *ChromatinAssay* class for single-cell chromatin data analysis. We extended the *Assay* class to add slots for the storage of genomic ranges, DNA motifs, genome build information, gene annotations, Tn5 insertion bias, positional enrichment information, genomic links, and linked on-disk data storage as fragment files.

The fragment file is a data format introduced by 10x Genomics for the storage of scATAC-seq data. Fragment files are defined as coordinate-sorted, block gzip-compressed (bgzip) and indexed browser-extensible data (BED) files with the following five columns: chromosome, start, end, cell barcode, PCR duplicate count. The start and end fields of the fragment file correspond to positions of the two Tn5 integration events that generated the sequenced DNA fragment. As the fragment file contains a deduplicated and near-complete representation of a single-cell chromatin experiment, and existing tools are established to efficiently retrieve subsets of a fragment file that overlap a given set of genomic regions [Li, 2011, Morgan et al., 2020], we utilized the fragment file format as the central disk-based data structure in the Signac framework, and is the only requirement for running a single-cell data analysis using Signac.

To facilitate the construction of a fragment file outside of running the 10x Genomics Cellranger software, we developed a Python package (Sinto) capable of generating the fragment file from a BAM file. This software is available on the Python Package Index (PyPI; https://pypi.org/project/sinto/) and GitHub (https://github.com/timoast/sinto).

### Quality control metrics

#### Nucleosome signal

Nucleosome signal was defined as the ratio of mononucleosomal (147-294 bp) to nucleosome-free (<147 bp) fragments sequenced for the cell. To compute the nucleosome signal per cell, we sampled the first *n* fragments from the fragment file, where *n* was the total number of cells in the dataset multiplied by 10,000. We then divide the number of mononucleosomal fragments per cell by the number of nucleosome-free fragments. This was implemented in the NucleosomeSignal function in Signac.

#### TSS enrichment

The transcriptional start site (TSS) enrichment score was originally defined by the ENCODE consortium [ENCODE Project Consortium, 2012] as a signal-to-noise metric for ATAC-seq experiments. The TSS score was defined as the mean number of Tn5 insertion events centered on the TSS sites (+/- 500 bp of the TSS) divided by the mean Tn5 insertion count at TSS-flanking regions, defined as +900 to +1000 and -900 to -1000 bp from the TSS. Calculation of the TSS enrichment score per-cell was implemented in the TSSEnrichment function in Signac.

### Dimension reduction

#### Latent semantic indexing

Latent semantic indexing (LSI) involves two steps. First, we compute the term-frequency (TF) inverse-document-frequency (IDF) matrix from the count matrix. Term frequency was defined as *TF* = *C*_*i j*_*/F*_*j*_ where *C*_*i j*_ was the total number of counts for peak *i* in cell *j* and *F*_*j*_ was the total number of counts for cell *j*. Inverse document frequency was defined as *IDF* = *N/n*_*i*_, where *N* was the total number of cells in the dataset and *n*_*i*_ was the total number of counts for peak *i* across all cells. The TF-IDF matrix was then computed as *TFIDF* = *log*(1 + (*TF* × *IDF*) ×10^4^). For comparison with previously-published LSI methods [Cusanovich et al., 2015], we also computed *IDF* as *IDF* = *log*(1 + *N/n*_*i*_) and subsequently TF-IDF as *TF* × *IDF*. This was implemented in the RunTFIDF function in Signac, with the “method” argument used to choose the TF-IDF method used. We decomposed the resulting TF-IDF matrix via truncated singular value decomposition (SVD) using the irlba R package [Baglama et al., 2019, Baglama and Reichel, 2005], implemented in the RunSVD function in Signac.

### UMAP

We performed UMAP using the RunUMAP function in the Seurat package (v3.2.0) using LSI components 2 to 40 for the PBMC dataset, components 2 to 50 for the CRC tumor dataset, and 2 to 100 for the BICCN mouse brain dataset. The first LSI component was excluded from each analysis as it typically captures sequencing depth (technical variation), and was highly correlated with the total number of counts for the cell. The RunUMAP function uses the uwot R package to compute two-dimensional UMAP coordinates [McInnes and Healy, 2018, Melville, 2020].

### Genome browser visualization

A common analysis task for single-cell chromatin data is genome browser-style data visualization for different groups of cells. Signac enables such visualizations with cells dynamically grouped into different pseudo-bulk tracks by reading Tn5 integration data from a position-indexed fragment file [Li, 2011]. To visualize pseudo-bulk accessibility tracks for different groups of cells, we constructed a sparse matrix of base-resolution Tn5 integration events, where each row was a cell and each column a DNA base in the requested region. We then grouped cells and computed the total accessibility at each site within each group, and scaled the total accessibility within each group by the total number of cells in the group and the average total counts for cells in each group to account for differences in overall chromatin signal and cell number between different groups of cells. We then smoothed the chromatin signal across small regions by computing a rolling window sum across the genomic region for each group of cells (using a window size of 100 bp by default). This was implemented in the CoveragePlot function in Signac. We also implemented an interactive version of the CoveragePlot function using the Shiny Gadgets framework in R [Chang et al., 2020] as the CoverageBrowser function in Signac. The interactive CoverageBrowser provides the same functionality as CoveragePlot, but additionally allows interactive navigation to different genomic regions and dynamic regrouping of cells.

One major advantage of genome browser-style visualizations is the ability to stack different data visualizations conveying different information as different browser tracks. We further built this concept into the CoveragePlot and CoverageBrowser functions in Signac by including the ability to plot additional tracks displaying gene expression information, gene annotations, peak coordinates, genomic links, genomic ranges, or the presence/absence of Tn5 integration events in individual cells as genomic “tile” plots.

### PBMC analysis

We downloaded data for human PBMCs processed using the 10x Genomics Multiome (ATAC + RNA) method from the 10x Genomics website (https://support.10xgenomics.com/single-cell-multiome-atac-gex/datasets/1.0.0/pbmc_granulocyte_sorted_10k).

#### Quality control and cell filtering

We computed the nucleosome signal score and TSS enrichment score for each cell as described above. We retained cells with a TSS enrichment score greater than 1, a nucleosome signal score less than 2, between 1,000 and 100,000 total ATAC-seq counts (based on the 10x Cellranger ATAC-seq count matrix), and between 1,000 and 25,000 total RNA counts.

#### Gene expression data preprocessing and cell annotation

We normalized the gene expression UMI count data using SCTransform [Hafemeister and Satija, 2019] and performed principal component analysis (PCA) on the SCTransform Pearson residual matrix using the RunPCA function in Seurat. We found the 20 nearest neighbors for each cell using the FindNeighbors function, with dims=1:50 to use the first 50 principal components, and annotated cell types in the PBMC dataset by label transfer from a publicly available multimodal PBMC reference dataset [Hao et al., 2020]. We identified anchor cells [Stuart et al., 2019] between the query and reference datasets using the FindTransferAnchors function in Seurat v4, with reference.reduction=‘spca’ to use a precomputed reference dimensional reduction object. We then computed cell type predictions for each cell in the query using the TransferData function in Seurat. As erythrocytes are not nucleated and the query PBMC dataset was derived from cell nuclei, we assigned a small number of cells that were incorrectly predicted as erythrocytes to the most common predicted class of those cells’ 20 nearest neighbors.

#### DNA accessibility data processing

ATAC-seq peaks in the PBMC dataset were identified using MACS2 [Zhang et al., 2008] with the following arguments:-g 2.7e9 -f BED –nomodel –extsize 200 –shift -100 –max-gap 50. We used the fragment file as input to the peak calling algorithm, as this contained the deduplicated Tn5 insertion sites for each cell. Peak calling was performed for each cell type using the CallPeaks function in Signac, with group.by=“celltype” to call peaks on each predicted cell type separately and combine the resulting peak calls across all cell types. We removed any peaks overlapping annotated genomic blacklist regions for the hg38 genome [Amemiya et al., 2019]. We quantified counts for the resulting peak set for each cell using the FeatureMatrix function in Signac.

Dimension reduction was performed on the DNA accessibility assay dataset using LSI and UMAP as described above. We performed graph-based clustering on the LSI components 2 to 40 using the Smart Local Moving algorithm (function FindNeighbors with dimensions=2:40 and reduction=“lsi” followed by FindClusters with algorithm=3 in Seurat v3.2.0) [Waltman and van Eck, 2013].

#### Joint data visualization

We computed a weighted nearest neighbor (WNN) graph for the DNA accessibility and gene expression assays using the FindMultiModalNeighbors function in Seurat v4, with reduction.list=list(“pca”, “lsi”) and dims.list=list(1:50, 2:40) to use the PCA dimension reduction with dimensions 1 to 50 for the gene expression assay and the LSI dimension reduction with dimensions 2 to 40 for the DNA accessibility assay [Hao et al., 2020]. This produced a neighbor graph encompassing information from both data modalities. We then computed a 2-dimensional UMAP visualization using this WNN graph, using the RunUMAP function in Seurat, with nn.name=“wknn” to use the multimodal WNN graph.

#### Differential accessibility

Differentially accessibility between features was computed using the FindMarkers function in Seurat v3.2.0, using the logistic regression method [Ntranos et al., 2019] with the total number of counts in each group of cells added as a latent variable in the logistic regression models (method=“LR”, latent.vars=“nCount_ATAC”). We classified peaks with an adjusted p-value (Bonferroni corrected) less than 0.05 as being differentially accessible between the cell groups.

#### Motif enrichment

A hypergeometric test was used to test for overrepresentation of each DNA motif in the set of differentially accessible peaks compared to a background set of peaks. We tested motifs present in the JASPAR database [Fornes et al., 2020] for human (species code 9606) by first identifying which peaks contained each motif using the motifmatchr R package [Schep, 2020]. We computed the GC content (percentage of G and C nucleotides) for each differentially accessible peak and sampled a background set of 40,000 peaks such that the background set was matched for overall GC content, accessibility, and peak width. This was performed using the FindMotifs function in Signac, with features.match=c(“GC.percent”, “count”, “sequence.length”).

#### Motif footprinting

We performed transcription factor motif footprinting following previously described methods [Corces et al., 2018]. To account for Tn5 sequence insertion bias, we first computed the observed Tn5 insertion frequency at each DNA hexamer using all Tn5 insertions on chromosome 1. This was done by extracting the base-resolution Tn5 insertion positions for each fragment mapped to chromosome 1, and extending the insertion coordinate 3 bp upstream and 2 bp downstream. We then extracted the DNA sequence corresponding to these coordinates using the getSeq function from the Biostrings R package [Pagès et al., 2020] and counted the frequency of each hexamer using the table function in R. We next computed the expected Tn5 hexamer insertion frequencies based on the frequency of each hexamer on chromosome 1. We counted the frequency of each hexamer using the oligonucleotideFrequency function in the Biostrings package with width=6 and names=“chr1”, using the hg38 genome via the BSgenome R package [Pagès, 2020]. Finally, we computed the Tn5 insertion bias as the observed Tn5 insertions divided by the expected insertions at each hexamer. This was performed using the InsertionBias function in Signac.

To perform motif footprinting, we first identified the coordinates of each instance of the motif to be footprinted using the matchMotifs function from the motifmatchr package with out=“positions” to return the genomic coordinates of each motif instance [Schep, 2020]. Motif coordinates were then resized to include the +/-250 bp sequence. The Tn5 insertion frequency was counted at each position in the region for each motif instance to produce a matrix containing the total observed Tn5 insertion events at each position relative to the motif center for each cell. We then found the expected Tn5 insertion frequency matrix by computing the hexamer frequency matrix, *M*. The hexamer frequency matrix *M* was defined as a matrix with *i* rows corresponding to *i* different DNA hexamers and *j* columns corresponding to *j* positions centered on the motif, and each entry *M*_*i j*_ corresponded to the hexamer count for hexamer *i* at position *j*. To find the expected Tn5 insertion frequency at each position relative to the motif given the Tn5 insertion bias (see above), we computed the matrix cross product between the hexamer frequency matrix *M* and the Tn5 insertion bias vector. Finally, the expected Tn5 insertion frequencies were normalized by dividing by the mean expected frequency in the 50 bp flanking regions (the regions 200 to 250 bp from the motif). To correct for Tn5 insertion bias we subtracted the expected Tn5 insertion frequencies from the observed Tn5 insertion frequencies at each position. This was performed using the Footprint function in Signac.

#### Peak-to-gene linkage

We estimated a linkage score for each peak-gene pair using linear regression models, based on recent work described in the SHARE-seq method [Ma et al., 2020]. For each gene, we computed the Pearson correlation coefficient *r* between the gene expression and the accessibility of each peak within 500 kb of the gene TSS. For each peak, we then computed a background set of expected correlation coefficients given properties of the peak by randomly sampling 200 peaks located on a different chromosome to the gene, matched for GC content, accessibility, and sequence length (MatchRegionStats function in Signac). We then computed the Pearson correlation between the expression of the gene and the set of background peaks. A z-score was computed for each peak as *z* = (*r − µ*)*/σ*, where *µ* was the background mean correlation coefficient and *σ* was the standard deviation of the background correlation coefficients for the peak. We computed a p-value for each peak using a one-sided z-test, and retained peak-gene links with a p-value < 0.05 and a Pearson correlation coefficient > 0.05 or < -0.05. This was performed using the LinkPeaks function in Signac.

#### Fine-mapped eQTL analysis

eQTL variants for whole blood that were fine-mapped using CAVIAR [Hormozdiari et al., 2014] were downloaded from the GTEx v8 website (https://storage.googleapis.com/gtex_analysis_v8/single_tissue_qtl_data/GTEx_v8_finemapping_CAVIAR.tar) [GTEx Consortium, 2020]. For each fine-mapped eQTL overlapping a peak that was linked to a gene in our analysis we counted the number of times the eQTL-associated gene was the same as the linked gene. Cases where multiple fine-mapped eQTLs associated with the same gene overlapped the same peak were treated as a single variant. To find the expected overlap based on random chance, we selected a set of peaks for each gene at random from the peaks within 500 kb of the gene TSS, with the number of peaks selected equal to the number of linked peaks for that gene. We then repeated the same eQTL overlap analysis using the randomized link set, as described above.

#### Count downsampling analysis

To test the impact of sequencing depth and assay sensitivity on the performance of different TF-IDF methods, we down-sampled the total number of counts per cell from 100% (the full dataset) down to 80%, 60%, 40%, and 20% using the downsampleMatrix function from the DropletUtils R package [Griffiths et al., 2018, Lun et al., 2019]. For each downsampling, we re-ran TF-IDF using the Signac RunTFIDF function with either method=1 or method=2 to compare TF-IDF methods, followed by SVD and UMAP using the RunSVD and RunUMAP functions in Signac and Seurat. For each downsampling, we estimated how well the data structure was preserved compared to the full dataset by computing the k-nearest neighbor purity. This was defined as the fraction of k neighbors (where k=100) for each cell *i* that belonged to the same annotated cell type as cell *i*, where cell types were predicted as described above using the gene expression assay. We computed nearest neighbors using LSI components 2 to 40, using the RANN R package [Arya et al., 2019].

### Colorectal cancer analysis

#### scATAC-seq data processing

We downloaded processed scATAC-seq counts from Zenodo (https://zenodo.org/record/3977808) and the fragment file from NCBI GEO (GSE148509) and computed the nucleosome signal and TSS enrichment score per-cell as described above, and retained cells with over 1,000 counts and less than 50,000 counts, less than 5% of counts in genomic blacklist regions, TSS enrichment score over 3, and a nucleosome signal score less than 4, and a mitochondrial genome sequencing depth of equal or greater than 10. We performed dimension reduction using LSI and UMAP as described above, and identified clusters using the Smart Local Moving algorithm using the FindClusters function in Seurat with resolution=0.5 and algorithm=3 [Waltman and van Eck, 2013].

#### Mitochondrial variant detection

Single-cell mitochondrial variant data processed using mgatk [Lareau et al., 2020] was downloaded from Zenodo (https://zenodo.org/record/3977808), read into R using the Signac function ReadMGATK, and used to create a Seurat assay. Informative mitochondrial variants were identified using the IdentifyVariants function, which computes the strand concordance in variant counts (Pearson correlation) and the variance-mean ratio (VMR) for each variant, as previously described [Lareau et al., 2020]. Informative mitochondrial variants were selected with a VMR > 0.01 and strand concordance >= 0.65, provided the variant was confidently detected in >=5 cells. We then computed per-cell mitochondrial allele frequencies for informative variants using the AlleleFreq function in Signac.

#### Clonal clustering

We identified cell clones by performing graph-based clustering on the square-root-transformed allele frequency matrix by first creating a neighbor graph using the FindNeighbors function in Seurat with annoy.metric=“cosine” to use the cosine distance to define nearest neighbors, and with k=10. We then performed community detection using the Smart Local Moving algorithm [Waltman and van Eck, 2013] using the shared nearest-neighbor graph computed using Seurat. This was implemented in the FindClonotypes function in Signac.

### BICCN analysis and benchmarking

#### Raw data processing

We downloaded FASTQ files for the BICCN dataset from NeMO (https://nemoarchive.org/) and mapped the reads to the mm10 genome using BWA-MEM [Li, 2013]. We created a fragment file from the aligned BAM file using sinto (https://github.com/timoast/sinto) and tabix [Li, 2011]. We then identified peaks for each brain region using Genrich (https://github.com/jsh58/Genrich) with the -j parameter for ATAC-seq data. We retained peaks with a score over 950 and unified peaks across brain regions using the reduce function in the GenomicRanges R package [Lawrence et al., 2013]. After reducing peaks, we removed outlier peaks larger than 3,000 bp in length. This resulted in a total of 315,334 peaks. Code to produce the BICCN fragment file and unified peak set is available at https://github.com/timoast/BICCN.

#### Cell detection and region quantification

The total number of fragments per cell barcode was computed using the CountFragments function in Signac. This function was implemented using C++ and the Rcpp R package [Eddel-buettel and Balamuta, 2018]. The total fragment counts for each barcode was then used to determine which barcodes correspond to cells and which correspond to background DNA. We retained all barcodes with greater than 1,500 total fragment counts for initial peak quantification. We then quantified the number of fragments overlapping each peak for each cell using the Signac FeatureMatrix function. This function was parallelized via the future R package, allowing the user to determine the parallelization strategy used [Bengtsson, 2020]. Signac also includes a convenience function (GenomeBinMatrix) to quantify signal in genomic bins tiling the entire genome or given chromosomes. Following quantification, we retained cells containing over 1,000 total counts in peaks and peaks open in over 100 total cells.

#### Quality control and dimension reduction

We computed the nucleosome signal as described above, with n=1e9 to sample the first billion fragments in the fragment file. We also computed the TSS enrichment score per cell as described above, using gene annotations for the mouse genome from the EnsDb.Mmusculus.v79 R package. We reduced dimensionality using LSI and UMAP as described above.

#### Cell downsampling analysis

To test the scalability of key steps in the Signac workflow, we downsampled the BICCN dataset from the full dataset (733,526 cells), down to 50,000 cells. We randomly sampled different cell numbers from the full dataset and used the FilterCells function in Signac to create downsampled fragment files containing only the cells sampled. We then ran Feature-Matrix to quantify the total counts per peak (for all 315,334 peaks) using 1, 2, 4, or 8 cores, as well as NucleosomeSignal, TSSEnrichment, RunTFIDF, and RunSVD on each downsampled dataset and recorded the total runtime for each step in triplicate. For the FeatureMatrix step, we also profiled the maximum resident memory using the GNU time command.

## Acknowledgements

This work was supported by the Chan Zuckerberg Initiative (EOSS-0000000082 and HCA-A-1704-01895 to RS), and the National Institutes of Health (DP2HG009623-01, RM1HG011014-01 and OT2OD026673-01 to RS). CAL was supported by a Stanford Science Fellowship. We are grateful to L. Ludwig for insightful conversations about mtDNA lineage tracing. We thank Bing Ren for assistance in accessing the BICCN mouse brain dataset. We thank the CRAN maintainers for their assistance in distributing the Signac R package, and members of the Satija lab for feedback on the manuscript.

## Author contributions

TS and AS developed the Signac package with guidance from RS. RS supervised the research. CAL developed the mitochondrial lineage tracing methods and analysis. TS wrote the manuscript with input from all authors. All authors read and approved the final manuscript.

---

**Figure S1.**
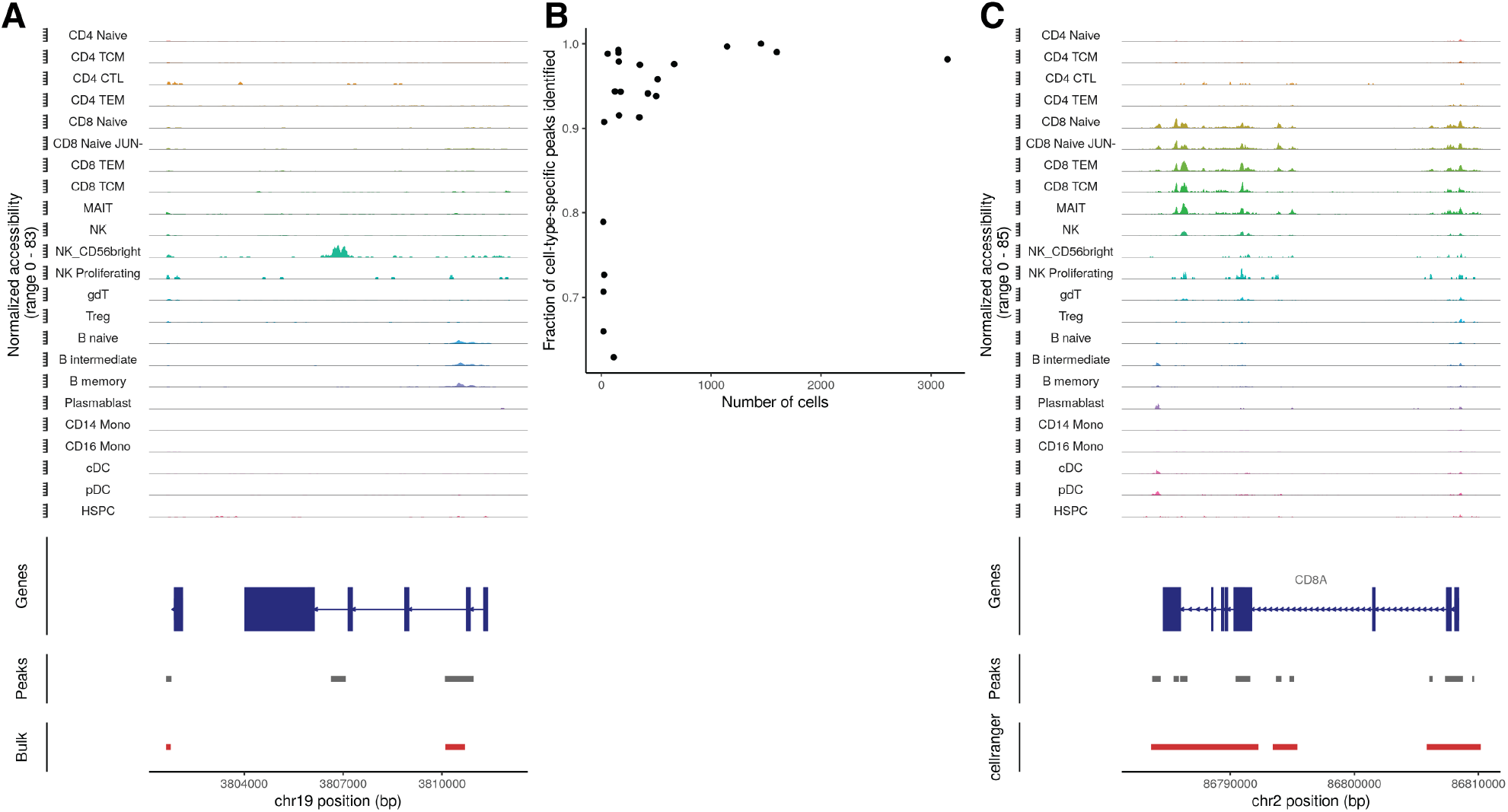
Comparison of peak calling methods for single-cell DNA accessibility data. (A) Comparison of cell-type-specific peak calling (grey) and bulk peak calling (red) using MACS2. (B) Fraction of cell-type-specific peaks for each cell type that were recovered when calling peaks on the bulk cell population. (C) Comparison of MACS2 (grey) and Cellranger (red) peak calls.

**Figure S2.**
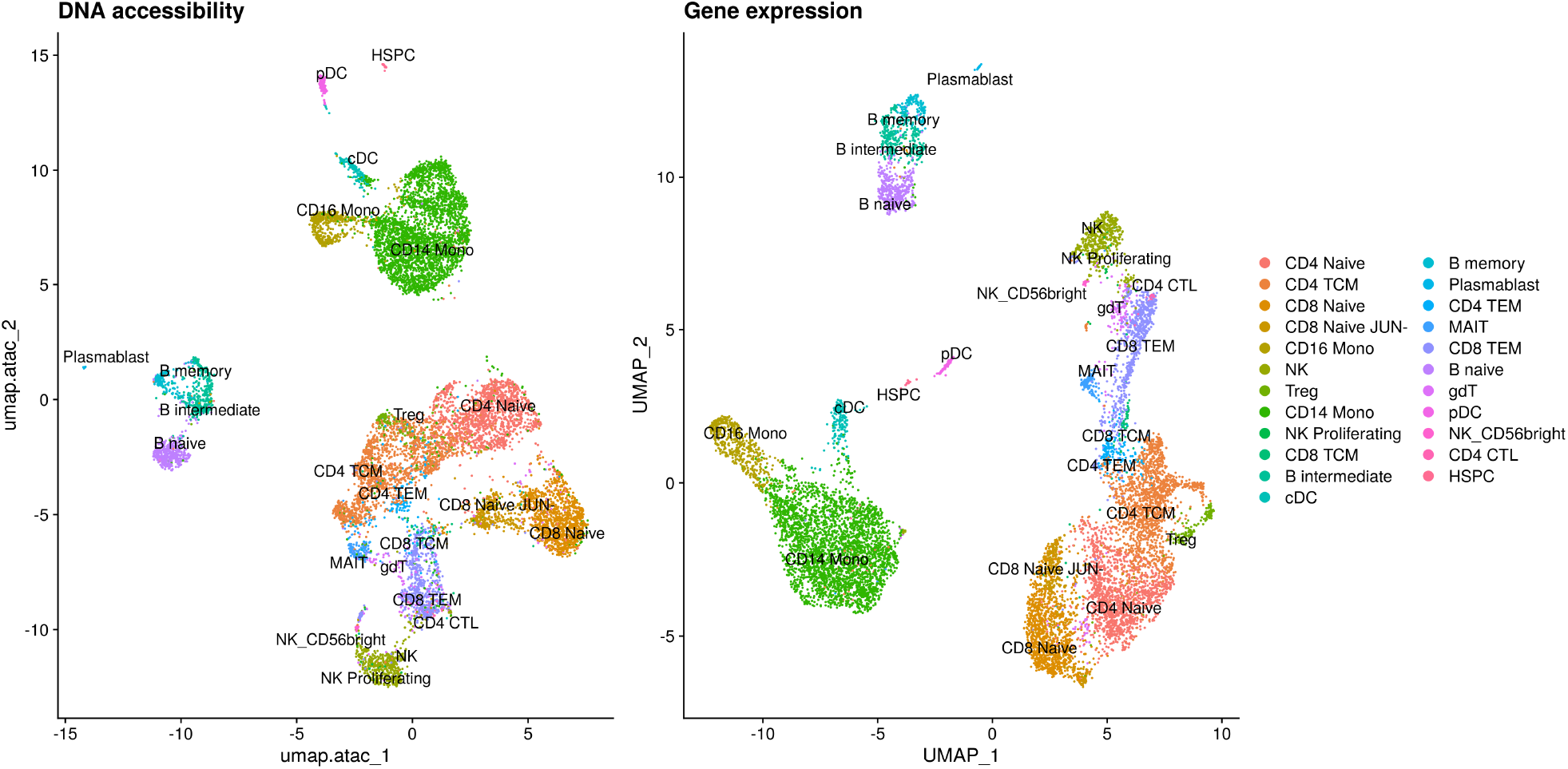
Reduced dimension representations of individual modalities. Separate UMAP plots constructed from the PBMC multiomic RNA assay or ATAC assay, with cells colored by predicted cell type.

## References

V. Agarwal and J. Shendure. Predicting mRNA abundance directly from genomic sequence using deep convolutional neural networks. Cell Rep., 31(7), May 2020. URL http://www.cell.com/article/S2211124720306161/abstract.

S. Ai, H. Xiong, C. C. Li, Y. Luo, Q. Shi, Y. Liu, X. Yu, C. Li, and A. He. Profiling chromatin states using single-cell itChIP-seq. Nat Cell Biol, 21(9):1164–1172, Sept. 2019. URL http://dx.doi.org/10.1038/s41556-019-0383-5.

H. M. Amemiya, A. Kundaje, and A. P. Boyle. The ENCODE blacklist: Identification of problematic regions of the genome. Sci. Rep., 9(1):9354, June 2019. URL https://www.nature.com/articles/s41598-019-45839-z.

S. Arora, M. Morgan, M. Carlson, and H. Pagès. Genome-InfoDb: Utilities for manipulating chromosome names, including modifying them to follow a particular naming style, 2020.

S. Arya, D. Mount, S. E. Kemp, and G. Jefferis. RANN: Fast nearest neighbour search (wraps ANN library) using L2 metric, 2019. URL “https://CRAN.R-project.org/package=RANN.

J. Baglama and L. Reichel. Augmented implicitly restarted lanczos bidiagonalization methods. SIAM J. Sci. Comput., 27(1):19–42, Jan. 2005. URL http:http://gateway.webofknowledge.com/gateway/Gateway.cgi?GWVersion=2&SrcAuth=mekentosj&SrcApp=Papers&DestLinkType=FullRecord&DestApp=WOS&KeyUT=000232354900002.

J. Baglama, L. Reichel, and B. W. Lewis. irlba: Fast truncated singular value decomposition and principal components analysis for large dense and sparse matrices, 2019. URL https://CRAN.R-project.org/package=irlba.

H. Bengtsson. future: Unified parallel and distributed processing in R for everyone, 2020. URL https://CRAN.R-project.org/package=future.

S. Brenner. Sequences and consequences. Philos. Trans. R. Soc. Lond. B Biol. Sci., 365(1537):207–212, Jan. 2010.URL http://dx.doi.org/10.1098/rstb.2009.0221.

J. D. Buenrostro, B. Wu, U. M. Litzenburger, D. Ruff, M. L. Gonzales, M. P. Snyder, H. Y. Chang, and W. J. Greenleaf. Single-cell chromatin accessibility reveals principles of regulatory variation. Nature, 523(7561):486–490, July 2015. URL http://dx.doi.org/10.1038/nature14590.

A. Butler, P. Hoffman, P. Smibert, E. Papalexi, and R. Satija. Integrating single-cell transcriptomic data across different conditions, technologies, and species. Nat. Biotechnol., Apr. 2018.URL http://dx.doi.org/10.1038/nbt.4096.

J. Cao, D. A. Cusanovich, V. Ramani, D. Aghamirzaie, H. A. Pliner, A. J. Hill, R. M. Daza, J. L. McFaline-Figueroa, J. S. Packer, L. Christiansen, F. J. Steemers, A. C. Adey, C. Trapnell, and J. Shendure. Joint profiling of chromatin accessibility and gene expression in thousands of single cells. Science, Aug. 2018.URL http://dx.doi.org/10.1126/science.aau0730.

J. Cao, M. Spielmann, X. Qiu, X. Huang, D. M. Ibrahim, A. J. Hill, F. Zhang, S. Mundlos, L. Christiansen, F. J. Steemers, C. Trapnell, and J. Shendure. The single-cell transcriptional landscape of mammalian organogenesis. Nature, page 1, Feb. 2019.URL https://www.nature.com/articles/s41586-019-0969-x.

B. Carter, W. L. Ku, J. Y. Kang, G. Hu, J. Perrie, Q. Tang, and K. Zhao. Mapping histone modifications in low cell number and single cells using antibody-guided chromatin tagmentation (ACT-seq). Nat. Commun., 10(1):3747, Aug. 2019.URL https://doi.org/10.1038/s41467-019-11559-1.

W. Chang, J. Cheng, J. J. Allaire, Y. Xie, and J. McPherson. shiny: Web application framework for R, 2020. URL https://CRAN.R-project.org/package=shiny.

S. Chen, B. B. Lake, and K. Zhang. High-throughput sequencing of the transcriptome and chromatin accessibility in the same cell. Nat. Biotechnol., Oct. 2019.URL https://doi.org/10.1038/s41587-019-0290-0.

S. J. Clark, R. Argelaguet, C.-A. Kapourani, T. M. Stubbs, H. J. Lee, C. Alda-Catalinas, F. Krueger, G. Sanguinetti, G. Kelsey, J. C. Marioni, O. Stegle, and W. Reik. scNMT-seq enables joint profiling of chromatin accessibility DNA methylation and transcription in single cells. Nat. Commun., 9(1):781, Feb. 2018.URL http://dx.doi.org/10.1038/s41467-018-03149-4.

M. R. Corces, J. M. Granja, S. Shams, B. H. Louie, J. A. Seoane, W. Zhou, T. C. Silva, C. Groeneveld, C. K. Wong, S. W. Cho, A. T. Satpathy, M. R. Mumbach, K. A. Hoadley, G. Robertson, N. C. Sheffield, I. Felau, M. A. A. Castro, P. Berman, L. M. Staudt, J. C. Zenklusen, P. W. Laird, Curtis, Cancer Genome Atlas Analysis Network, W. J. Greenleaf, and H. Y. Chang. The chromatin accessibility landscape of primary human cancers. Science, 362(6413), Oct. 2018.URL http://dx.doi.org/10.1126/science.aav1898.

A. Cusanovich, R. Daza, A. Adey, H. A. Pliner, L. Christiansen, K. L. Gunderson, F. J. Steemers, C. Trapnell, and J. Shendure. Multiplex single cell profiling of chromatin accessibility by combinatorial cellular indexing. Science, 348(6237):910–914, May 2015. URL http://dx.doi.org/10.1126/science.aab1601.

A. Danese, M. L. Richter, D. S. Fischer, F. J. Theis, and M. Colomé-Tatché. EpiScanpy: integrated single-cell epigenomic analysis. May 2019. URL https://www.biorxiv.org/content/10.1101/648097v1.

S. Deerwester, S. T. Dumais, G. W. Furnas, T. K. Landauer, and R. Harshman. Indexing by latent semantic analysis. Journal of the American Society for Information Science, 41(6):391–407, Sept. 1990.URL https://doi.org/10.1002/(SICI)1097-4571(199009)41:6<391::AID-ASI1>3.0.CO;2-9.

D. Eddelbuettel and J. J. Balamuta. Extending R with c++: A brief introduction to rcpp. Am. Stat., 72(1):28–36, Jan. 2018.URL https://doi.org/10.1080/00031305.2017.1375990.

ENCODE Project Consortium. An integrated encyclopedia of DNA elements in the human genome. Nature, 489(7414): 57–74, Sept. 2012.URL http://www.nature.com/doifinder/10.1038/nature11247.

R. Fang, S. Preissl, X. Hou, J. Lucero, X. Wang, A. Motamedi, A. K. Shiau, E. A. Mukamel, Y. Zhang, M. Margarita Behrens, J. Ecker, and B. Ren. Fast and accurate clustering of single cell epigenomes reveals Cis-Regulatory elements in rare cell types. Apr. 2019.URL https://www.biorxiv.org/content/10.1101/615179v1.

E. Fiskin, C. A. Lareau, G. Eraslan, L. S. Ludwig, and A. Regev. Single-cell multimodal profiling of proteins and chromatin accessibility using PHAGE-ATAC. Oct. 2020.URL https://www.biorxiv.org/content/10.1101/2020.10.01.322420v1.

O. Fornes, J. A. Castro-Mondragon, A. Khan, R. van der Lee, X. Zhang, P. A. Richmond, B. P. Modi, S. Correard, M. Gheorghe, D. Baranašić, W. Santana-Garcia, G. Tan, J. Chèneby, B. Ballester, F. Parcy, A. Sandelin, B. Lenhard,W. W. Wasserman, and A. Mathelier. JASPAR 2020: update of the open-access database of transcription factor binding profiles. Nucleic Acids Res., 48(D1):pD87–D92, Jan. 2020.URL http://dx.doi.org/10.1093/nar/gkz1001.

R. C. Gentleman, V. J. Carey, D. M. Bates, B. Bolstad, M. Dettling, S. Dudoit, B. Ellis, L. Gautier, Y. Ge, J. Gentry, K. Hornik, T. Hothorn, W. Huber, S. Iacus, R. Irizarry, F. Leisch, C. Li, M. Maechler, A. J. Rossini, G. Sawitzki, C. Smith, G. Smyth, L. Tierney, J. Y. H. Yang, and J. Zhang. Bioconductor: open software development for computational biology and bioinformatics. Genome Biol., 5(10):R80, Sept. 2004.URL http://dx.doi.org/10.1186/gb-2004-5-10-r80.

J. M. Granja, M. Ryan Corces, S. E. Pierce, S. Tansu Bagdatli, H. Choudhry, H. Chang, and W. Greenleaf. ArchR: An integrative and scalable software package for single-cell chromatin accessibility analysis. Apr. 2020.URL https://www.biorxiv.org/content/10.1101/2020.04.28.066498v1.

J. A. Griffiths, A. C. Richard, K. Bach, A. T. L. Lun, and J. C. Marioni. Detection and removal of barcode swapping in single-cell RNA-seq data. Nat. Commun., 9(1):2667, July 2018. URL http://dx.doi.org/10.1038/s41467-018-05083-x.

GTEx Consortium. The GTEx consortium atlas of genetic regulatory effects across human tissues. Science, 369(6509):1318–1330, Sept. 2020.URL http://dx.doi.org/10.1126/science.aaz1776.

C. Hafemeister and R. Satija. Normalization and variance stabilization of single-cell RNA-seq data using regularized negative binomial regression. Genome Biol., 20(1):296, Dec. 2019.URL http://dx.doi.org/10.1186/s13059-019-1874-1.

Y. Hao, S. Hao, E. Andersen-Nissen, W. M. Mauck, S. Zheng, A. Butler, M. J. Lee, A. J. Wilk, C. Darby, M. Zagar, P. Hoffman, M. Stoeckius, E. Papalexi, E. P. Mimitou, J. Jain, A. Srivastava, T. Stuart, L. B. Fleming, B. Yeung, A. J. Rogers, J. M. McElrath, C. A. Blish, R. Gottardo, P. Smibert, and R. Satija. Integrated analysis of multimodal single-cell data. Oct. 2020.URL https://www.biorxiv.org/content/10.1101/2020.10.12.335331v1.

F. Hormozdiari, E. Kostem, E. Y. Kang, B. Pasaniuc, and E. Eskin. Identifying causal variants at loci with multiple signals of association. Genetics, 198(2):497–508, Oct. 2014.URL http://dx.doi.org/10.1534/genetics.114.167908.

W. Huber, V. J. Carey, R. Gentleman, S. Anders, M. Carlson, B. S. Carvalho, H. C. Bravo, S. Davis, L. Gatto, T. Girke, R. Gottardo, F. Hahne, K. D. Hansen, R. A. Irizarry, M. Lawrence, M. I. Love, J. MacDonald, V. Obenchain, A. K. Oleś, H. Pagès, A. Reyes, P. Shannon, G. K. Smyth, D. Tenenbaum, L. Waldron, and M. Morgan. Orchestrating high-throughput genomic analysis with bioconductor. Nat. Methods, 12(2):115–121, Feb. 2015.URL http://dx.doi.org/10.1038/nmeth.3252.

H. S. Kaya-Okur, S. J. Wu, C. A. Codomo, E. S. Pledger, T. D. Bryson, J. G. Henikoff, K. Ahmad, and S. Henikoff. CUT&Tag for efficient epigenomic profiling of small samples and single cells. Nat. Commun., 10(1):1930, Apr. 2019.URL https://doi.org/10.1038/s41467-019-09982-5.

D. R. Kelley, Y. A. Reshef, M. Bileschi, D. Belanger, C. Y. McLean, and J. Snoek. Sequential regulatory activity prediction across chromosomes with convolutional neural networks. Genome Res., 28(5):739–750, May 2018. URL http://dx.doi.org/10.1101/gr.227819.117.

I. Korsunsky, N. Millard, J. Fan, K. Slowikowski, F. Zhang, K. Wei, Y. Baglaenko, M. Brenner, P.-R. Loh, and S. Raychaudhuri. Fast, sensitive and accurate integration of single-cell data with harmony. Nat. Methods, Nov. 2019.URL https://doi.org/10.1038/s41592-019-0619-0.

C. A. Lareau, F. M. Duarte, J. G. Chew, V. K. Kartha, Z. D. Burkett, A. S. Kohlway, D. Pokholok, M. J. Aryee, F. J. Steemers, R. Lebofsky, and J. D. Buenrostro. Droplet-based combinatorial indexing for massive-scale single-cell chromatin accessibility. Nat Biotechnol, June 2019. URL http://dx.doi.org/10.1038/s41587-019-0147-6.

C. A. Lareau, L. S. Ludwig, C. Muus, S. H. Gohil, T. Zhao, Z. Chiang, K. Pelka, J. M. Verboon, W. Luo, E. Christian, D. Rosebrock, G. Getz, G. M. Boland, F. Chen, J. D. Buenrostro, N. Hacohen, C. J. Wu, M. J. Aryee, A. Regev, and V. G. Sankaran. Massively parallel single-cell mitochondrial DNA genotyping and chromatin profiling. Nat. Biotechnol., pages 1–12, Aug. 2020.URL http://dx.doi.org/10.1038/s41587-020-0645-6.

M. Lawrence, W. Huber, H. Pagès, P. Aboyoun, M. Carlson, R. Gentleman, M. T. Morgan, and V. J. Carey. Software for computing and annotating genomic ranges. PLoS Comput. Biol., 9(8):e1003118, Aug. 2013.URL http://dx.doi.org/10.1371/journal.pcbi.1003118.

H. Li. Tabix: fast retrieval of sequence features from generic TAB-delimited files. Bioinformatics, 27(5):718–719, Mar. 2011.URL http://dx.doi.org/10.1093/bioinformatics/btq671.

H. Li. Aligning sequence reads, clone sequences and assembly contigs with BWA-MEM. arXiv.org, Jan. 2013.URL http://arxiv.org/abs/1303.3997.

Y. Li, S. Preissl, X. Hou, Z. Zhang, K. Zhang, R. Fang, Y. Qiu, O. Poirion, B. Li, Y. Yan, H. Liu, X. Wang, J. Y. Han, J. Lucero, S. Kuan, D. Gorkin, M. Nunn, E. A. Mukamel, M. Margarita Behrens, J. R. Ecker, and B. Ren. An atlas of gene regulatory elements in adult mouse cerebrum. May 2020. URL https://www.biorxiv.org/content/10.1101/2020.05.10.087585v1?ct=.

L. Liu, C. Liu, A. Quintero, L. Wu, Y. Yuan, M. Wang, M. Cheng, L. Leng, L. Xu, G. Dong, R. Li, Y. Liu, X. Wei, J. Xu, X. Chen, H. Lu, D. Chen, Q. Wang, Q. Zhou, X. Lin, G. Li, S. Liu, Q. Wang, H. Wang, J. L. Fink, Z. Gao, X. Liu, Y. Hou, S. Zhu, H. Yang, Y. Ye, G. Lin, F. Chen, C. Herrmann, R. Eils, Z. Shang, and X. Xu. Deconvolution of single-cell multi-omics layers reveals regulatory heterogeneity. Nat. Commun., 10(1):470, Jan. 2019.URL http://dx.doi.org/10.1038/s41467-018-08205-7.

L. S. Ludwig, C. A. Lareau, J. C. Ulirsch, E. Christian, C. Muus, L. H. Li, K. Pelka, W. Ge, Y. Oren, A. Brack, T. Law, C. Rodman, J. H. Chen, G. M. Boland, N. Hacohen, O. Rozenblatt-Rosen, M. J. Aryee, J. D. Buenrostro, A. Regev, and V. G. Sankaran. Lineage tracing in humans enabled by mitochondrial mutations and Single-Cell genomics. Cell, 0(0), Feb. 2019.URL http://www.cell.com/article/S0092867419300558/abstract.

A. T. L. Lun, S. Riesenfeld, T. Andrews, T. P. Dao, T. Gomes, participants in the 1 st Human Cell Atlas Jamboree, and J. C. Marioni. EmptyDrops: distinguishing cells from empty droplets in droplet-based single-cell RNA sequencing data. Genome Biol., 20(1):63, Mar. 2019.URL http://dx.doi.org/10.1186/s13059-019-1662-y.

C. Luo, A. Rivkin, J. Zhou, J. P. Sandoval, L. Kurihara, J. Lucero, R. Castanon, J. R. Nery, A. Pinto-Duarte, B. Bui, C. Fitzpatrick, C. O’Connor, S. Ruga, M. E. Van Eden, D. A. Davis, D. C. Mash, M. M. Behrens, and J. R. Ecker. Robust single-cell DNA methylome profiling with snmc-seq2. Nat. Commun., 9(1):3824, Sept. 2018.URL http://dx.doi.org/10.1038/s41467-018-06355-2.

S. Ma, B. Zhang, L. M. LaFave, A. S. Earl, Z. Chiang, Y. Hu, J. Ding, A. Brack, V. K. Kartha, T. Tay, T. Law, C. Lareau, Y.-C. Hsu, A. Regev, and J. D. Buenrostro. Chromatin potential identified by shared Single-Cell profiling of RNA and chromatin. Cell, 0(0), Oct. 2020.URL http://www.cell.com/article/S0092867420312538/abstract.

L. McInnes and J. Healy. UMAP: Uniform manifold approximation and projection for dimension reduction. Feb. 2018.URL http://arxiv.org/abs/1802.03426.

J. Melville. uwot: The uniform manifold approximation and projection (UMAP) method for dimensionality reduction, 2020. URL https://CRAN.R-project.org/package=uwot.

E. P. Mimitou, C. A. Lareau, K. Y. Chen, A. L. Zorzetto-Fernandes, Y. Takeshima, W. Luo, T.-S. Huang, B. Yeung, P. I. Thakore, J. B. Wing, K. L. Nazor, S. Sakaguchi, L. S. Ludwig, V. G. Sankaran, A. Regev, and P. Smibert. Scalable, multimodal profiling of chromatin accessibility and protein levels in single cells. Sept. 2020.URL https://www.biorxiv.org/content/10.1101/2020.09.08.286914v1.

M. Morgan, H. Pagès, V. Obenchain, and N. Hayden. Rsamtools: Binary alignment (bam), fasta, variant call (bcf), and tabix file import, 2020. URL http://bioconductor.org/packages/Rsamtools.

V. Ntranos, L. Yi, P. Melsted, and L. Pachter. A discriminative learning approach to differential expression analysis for single-cell RNA-seq. Nat. Methods, 16(2):163–166, Feb. 2019.URL http://dx.doi.org/10.1038/s41592-018-0303-9.

H. Pagès. BSgenome: Software infrastructure for efficient representation of full genomes and their SNPs, 2020.

H. Pagès, P. Aboyoun, R. Gentleman, and S. DebRoy. Biostrings: Efficient manipulation of biological strings, 2020.

E. L. Pearce, A. C. Mullen, G. A. Martins, C. M. Krawczyk, A. S. Hutchins, V. P. Zediak, M. Banica, C. B. DiCioccio, D. A. Gross, C.-A. Mao, H. Shen, N. Cereb, S. Y. Yang, T. Lindsten, J. Rossant, C. A. Hunter, and S. L. Reiner. Control of effector CD8+ T cell function by the transcription factor eomesodermin. Science, 302(5647):1041–1043, Nov. 2003.URL http://dx.doi.org/10.1126/science.1090148.

S. E. Pierce, J. M. Granja, and W. J. Greenleaf. High-throughput single-cell chromatin accessibility CRISPR screens enable unbiased identification of regulatory networks in cancer. Nov. 2020.URL https://www.biorxiv.org/content/10.1101/2020.11.02.364265v1?rss=1.

H. A. Pliner, J. S. Packer, J. L. McFaline-Figueroa, D. A. Cusanovich, R. M. Daza, D. Aghamirzaie, S. Srivatsan, X. Qiu, D. Jackson, A. Minkina, A. C. Adey, F. J. Steemers, J. Shendure, and C. Trapnell. Cicero predicts cis-regulatory DNA interactions from Single-Cell chromatin accessibility data. Mol. Cell, 71(5):858–871.e8, Sept. 2018.URL http://dx.doi.org/10.1016/j.molcel.2018.06.044.

A. J. Rubin, K. R. Parker, A. T. Satpathy, Y. Qi, B. Wu, A. J. Ong, M. R. Mumbach, A. L. Ji, D. S. Kim, S. W. Cho, B. J. Zarnegar, W. J. Greenleaf, H. Y. Chang, and P. A. Khavari. Coupled Single-Cell CRISPR screening and epigenomic profiling reveals causal gene regulatory networks. Cell, 176(1-2):361–376.e17, Jan. 2019.URL http://dx.doi.org/10.1016/j.cell.2018.11.022.

R. Satija, J. A. Farrell, D. Gennert, A. F. Schier, and A. Regev. Spatial reconstruction of single-cell gene expression data. Nat. Biotechnol., 33(5):495–502, May 2015. URL http://www.nature.com/doifinder/10.1038/nbt.3192.

A. T. Satpathy, J. M. Granja, K. E. Yost, Y. Qi, F. Meschi, G. P. McDermott, B. N. Olsen, M. R. Mumbach, S. E. Pierce, M. Ryan Corces, P. Shah, J. C. Bell, D. Jhutty, C. M. Nemec, J. Wang, L. Wang, Y. Yin, P. G. Giresi, A. L. S. Chang, G. X. Y. Zheng, W. J. Greenleaf, and H. Y. Chang. Massively parallel single-cell chromatin landscapes of human immune cell development and intratumoral T cell exhaustion. Nat. Biotechnol., 37(8):925–936, Aug. 2019.URL https://www.nature.com/articles/s41587-019-0206-z.

A. Schep. motifmatchr: Fast motif matching in R, 2020.

A. N. Schep, B. Wu, J. D. Buenrostro, and W. J. Greenleaf. chromVAR: inferring transcription-factor-associated accessibility from single-cell epigenomic data. Nat. Methods, 14 (10):975–978, Oct. 2017.URL http://dx.doi.org/10.1038/nmeth.4401.

S. A. Smallwood, H. J. Lee, C. Angermueller, F. Krueger, H. Saadeh, J. Peat, S. R. Andrews, O. Stegle, W. Reik, and G. Kelsey. Single-cell genome-wide bisulfite sequencing for assessing epigenetic heterogeneity. Nat. Methods, 11(8):817–820, July 2014. URL http://www.nature.com/doifinder/10.1038/nmeth.3035.

T. Stuart and R. Satija. Integrative single-cell analysis. Nat. Rev. Genet., Jan. 2019.URL http://dx.doi.org/10.1038/s41576-019-0093-7.

T. Stuart, A. Butler, P. Hoffman, C. Hafemeister, E. Papalexi, W. M. Mauck, 3rd, Y. Hao, M. Stoeckius, P. Smibert, and R. Satija. Comprehensive integration of Single-Cell data. Cell, 177(7):1888–1902.e21, June 2019. URL http://dx.doi.org/10.1016/j.cell.2019.05.031.

E. Swanson, C. Lord, J. Reading, A. T. Heubeck, A. K. Savage, R. Green, T. R. Torgerson, T. F. Bumol, L. T. Graybuck, and P. J. Skene. Integrated single cell analysis of chromatin accessibility and cell surface markers. Sept. 2020.URL https://www.biorxiv.org/content/10.1101/2020.09.04.283887v1.

C. A. Thornton, R. M. Mulqueen, K. A. Torkenczy, E. G. Lowenstein, A. J. Fields, F. J. Steemers, K. M. Wright, and A. C. Adey. Spatially-mapped single-cell chromatin accessibility. Oct. 2019.URL https://www.biorxiv.org/content/10.1101/815720v1.

L. Waltman and N. J. van Eck. A smart local moving algorithm for large-scale modularity-based community detection. Eur. Phys. J. B, 86(11):471, Nov. 2013.URL https://doi.org/10.1140/epjb/e2013-40829-0.

H. Wickham. ggplot2: Elegant graphics for data analysis, 2016. URL https://ggplot2.tidyverse.org.

Q. R. Xing, C. A. E. Farran, Y. Y. Zeng, Y. Yi, T. Warrier, P. Gautam, J. J. Collins, J. Xu, P. Dröge, C.-G. Koh, H. Li, L.-F. Zhang, and Y.-H. Loh. Parallel bimodal single-cell sequencing of transcriptome and chromatin accessibility. Genome Res., 30(7):1027–1039, July 2020. URL http://dx.doi.org/10.1101/gr.257840.119.

J. Xu, K. Nuno, U. M. Litzenburger, Y. Qi, M. R. Corces, R. Majeti, and H. Y. Chang. Single-cell lineage tracing by endogenous mutations enriched in transposase accessible mitochondrial DNA. Elife, 8, Apr. 2019.URL http://dx.doi.org/10.7554/eLife.45105.

Y. Zhang, T. Liu, C. A. Meyer, J. Eeckhoute, D. S. Johnson, B. E. Bernstein, C. Nussbaum, R. M. Myers, M. Brown, W. Li, and X. S. Shirley. Model-based analysis of ChIP-Seq (MACS). Genome Biol., 9(9):R137, Sept. 2008.URL http://genomebiology.biomedcentral.com/articles/10.1186/gb-2008-9-9-r137.

C. Zhu, M. Yu, H. Huang, I. Juric, A. Abnousi, R. Hu, J. Lucero, M. M. Behrens, M. Hu, and B. Ren. An ultra high-throughput method for single-cell joint analysis of open chromatin and transcriptome. Nat. Struct. Mol. Biol., 26(11):1063–1070, Nov. 2019.URL http://dx.doi.org/10.1038/s41594-019-0323-x.

